# SmartHisto: Bayesian Active Learning for Histology Images

**DOI:** 10.1101/2025.10.13.681994

**Authors:** Sriram Vijendran, Bailey Arruda, Tavis K. Anderson, Oliver Eulenstein

## Abstract

Accurate and efficient characterization of biological images is crucial for advancing systems biology and medical research. Recent advancements in deep learning and image processing have enabled neural network models to rapidly accelerate image analysis by utilizing large expert-annotated datasets. However, in histopathology, the size of whole-slide images makes expert annotation expensive, limiting the acquisition of sufficiently large annotated datasets and posing a major challenge for developing automated, AI-driven image analysis pipelines. To address this limitation, we propose a novel active learning-based framework to train image segmentation models interactively. Our approach employs a Bayesian neural network to identify informative regions in unlabeled images rather than entire images, making expert labeling more cost-effective. We validate our framework on multiple benchmark datasets spanning different staining techniques and magnifications, demonstrating substantial reductions in annotation effort. Notably, our method achieves a mean IoU of 0.75, significantly outperforming competing approaches, which average 0.60.

**Author summary:** Histopathology is fundamental to investigating tissue and immune responses, host-pathogen interactions, and disease mechanisms. However, histopathology is highly resource-intensive and requires specialized training, dramatically increasing the costs of annotating whole-slide images and, consequently, the expenses of large-scale studies involving numerous labs and specialists. We developed a computational tool to overcome these challenges, implementing a robust uncertainty-based sampling algorithm in conjunction with a next-generation Bayesian Convolutional Neural Network. This algorithm can be used for hypothesis testing and discovery by reducing the reliance on large, precisely annotated training datasets required in automated image analysis pipelines. The base model, when trained to identify lung tissue types using a small set of annotated images, outperforms state-of-the-art models and can efficiently annotate thousands of images much more quickly than a human. Models trained by the proposed algorithm will serve as a standardized approach for pathologists and disease researchers to train automated image segmentation pipelines for large-scale histopathology.

## Introduction

Semantic image segmentation, or pixel-level classification, is the task of partitioning an image into distinct regions or segments, each representing different objects or areas of interest. In recent years, deep neural networks trained on large, expert-annotated datasets have significantly advanced this field, achieving near-human performance as dataset sizes have increased [1,2]. Semantic image segmentation has been successfully applied to domains such as automated driving [3], intelligent medical technology [4–6], image search engines [7,8], industrial inspection [9–11], and augmented reality [1].

Despite its success, semantic image segmentation has seen limited application in histopathology, a critical discipline for understanding host-pathogen interactions and immune responses. Histopathology is a specialized technique that involves microscopic examination of tissue samples to investigate and analyze disease mechanisms. However, histologic evaluation requires years of specialized training and costly equipment. Consequently, only a fraction of collected specimens is analyzed, with vast amounts of archival material remaining underutilized. Moreover, the high costs associated with image annotation create bias and accuracy challenges [12,13], limiting the effectiveness of machine learning-based analysis pipelines.

Active Learning (AL) offers a promising approach for facilitating access and utilization of archived images to train AI models, where informative subsets are sampled from a pool of unlabeled data for annotation by human experts, with the goal of selecting the fewest samples to learn a robust neural network model [14,15]. AL heuristics quantify the information within a sample by prediction uncertainty [16–22]. AL has produced highly accurate models in classification tasks with data scarcity due to high data annotation costs [23,24].

However, AL is challenging to apply in image segmentation [25–28] due to two key factors: 1) sample selection and 2) uncertainty quantification. Individual pixels in an image must be classified and may carry varying degrees of uncertainty [26,28]. Images containing only a few pixels with high uncertainty can skew sample selection and confound sample ranking, resulting in biased and subpoptimal models. Furthermore, classical neural network models, also called point-estimate models, cannot differentiate between uncertainty due to ambiguity in the data-generating process (aleatoric uncertainty) or lack of information on the data-generating process (epistemic uncertainty). Epistemic uncertainty arises due to a lack of understanding and can be reduced by acquiring more data. In comparison, aleatoric uncertainty is due to inherent noise and is hence irreducible.

To address these challenges, we propose SmartHisto, a novel sample selection heuristic designed for the semantic segmentation of histology whole-slide images. SmartHisto trains a Bayesian Neural Network (BNN) [29,30] instead of a point-estimate model to improve uncertainty quantification and identify the individual structures that are present in each image.

### Related Work

AL is a methodology for achieving high model performance with minimal labeled data by iteratively selecting the most informative samples for annotation. Computationally, AL is a set selection problem, where in each iteration, a subset of all available data points should be selected to maximize some measure of information gain. Such informative samples are sequentially selected, annotated by an oracle, and then used to train a machine learning model, such as a Neural Network. AL has been tackled using uncertainty-based [25,28,31,32], diversity-based [33], and expected model change-based approaches [27,34]. These three methods have previously been integrated within several deep learning models.

While AL has been used extensively in classification [23,24] where the model attempts to assign a single label for a sample, application of AL in image segmentation using classical point-estimate model [25–28] remains challenging. Unlike point-estimate models, BNNs [29,30] provide a more robust measure of uncertainty that differentiates between aleatoric and epistemic uncertainty. BNNs sample model parameters from distributions rather than use individual fixed values; BNNs can better identify and quantify sources of uncertainty than classical Neural Networks [35]. The capability to distinguish novel from familiar information makes the application of BNNs in AL frameworks more attractive for complex tasks such as semantic image segmentation.

### Our Contribution

This article proposes SmartHisto, a novel sample selection heuristic for semantic image segmentation using BNNs. SmartHisto differentiates between *aleatoric* and *epistemic* uncertainty and employs superpixeling [36] to cluster pixels into superpixels based on the measured uncertainty. A novel divergence-based uncertainty metric is then computed to rank samples according to their superpixels, optimizing sample selection and streamlining label acquisition.

Our experimental studies on a standardized benchmark dataset and a specially curated lung pathology dataset indicate that our proposed sampler produces superior models to traditional samplers while using fewer training samples. Finally, our comparison study indicates that models trained using SmartHisto outperform current state-of-the-art (SOTA) image segmentation models, such as QBC [37] and DEAL [38], without compromising memory efficiency.

The structure of this paper is as follows. The Materials and methods section formalizes the architecture of the BNN used for image segmentation in conjunction with algorithms to quantify uncertainty and select samples for annotation. The Results section presents the metrics used to evaluate the proposed method, an experimental setup with publicly available benchmarking datasets, and a comparison study that contrasts our approach with current state-of-the-art AL algorithms. Finally, the Discussion section concludes the article with future directions and potential applications.

## Materials and methods

The proposed approach consists of two significant state-of-the-art advances: model initialization, referred to as *learner*, and uncertainty estimation. The BNN structure is described in Section Bayesian Neural Network Architecture, where the parameters of a regular point-estimate model are replaced with distributions from which parameters are sampled during inference. To train a BNN for semantic image segmentation, a loss function that capitalizes on the dual-parameter initialization is formalized in Section Evaluating Loss in a Bayesian Neural Network. Next, a heuristic to approximate the pixelwise uncertainty is introduced in Section Estimating Pixelwise Uncertainty. Finally, to address the challenge of annotating individual pixels, a superpixeling method is described in Section Selecting Regions for Annotation.

### Bayesian Neural Network Architecture

The target learner architecture used in this study is based on the UNet [39] architecture. This approach replaces the regular point estimate initialization used by AL paradigms with parameters, each sampled from Gaussian distributions. The approach uses a smaller version of the traditional UNet architecture with weights drawn from independent Gaussian distributions.

BNNs replace the deterministic network’s weight parameters with distributions over these parameters. Instead of optimizing the network weights directly, our approach optimizes the network using the marginalized weight distribution. The BNN is denoted byℳ, and the model prediction for some input image *x* is given by ℳ (*x*). The model likelihood is defined as *p*(*y*|ℳ (*x*)). Given a dataset with *m* samples with input features, *X*, and target labels, *Y*, Bayesian inference computes the posterior distribution over the weights given by *p*(*w* |*X, Y*).

A distribution-driven initialization results in a learner that behaves as an ensemble of models trained with the same dataset, i.e., an ensemble of models with identical priors but different posterior weight distributions. This form of parameter initialization results in a more reliable measure of uncertainty as the ensemble of models are highly similar; In this setting, computing the uncertainty of a prediction is equivalent to the variance in prediction per pixel.

The target learner is trained using Bayes by Backprop [40], where each parameter 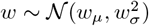 is given by a pair (*w*_*µ*_, *w*_*ρ*_), where *w*_*σ*_ is computed from *w*_*ρ*_ as given below.

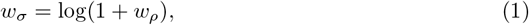

Following the weight initialization, the weights and biases of the target learner are given by:

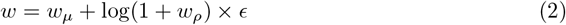

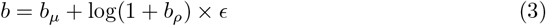

where *ϵ* is added Gaussian noise i.e, *ϵ* ~*𝒩* (1, 0).

The initialization for all convolution layers follows the methodology described in [41], where a single convolution layer is equivalent to two sequential convolution operations.

The final output layer uses a sigmoid activation to compute individual class probabilities for each pixel. This approach allows the learner to better estimate the uncertainty of a superpixel, thereby identifying samples in the unlabeled pool of images from which to train. The proposed formulation of the target learner allows efficient characterization of the different types of uncertainty, discussed further in Section Estimating Pixelwise Uncertainty.

### Evaluating Loss in a Bayesian Neural Network

Various methods exist to compute the loss in semantic segmentation. Common approaches include Mean Intersection over Union (mIoU) [42,43], Jaccard Similarity [44,45], and Tversky Loss (TI) [46,47]. We use Dice Binary Cross entropy (DiceBCE) [44] to evaluate the quality of the predicted mask, whose formulation accommodates the hierarchical classes frequently seen in histopathology. DiceBCE is computed by adding the binary cross entropy loss with the *Dice Loss*, which is calculated as:

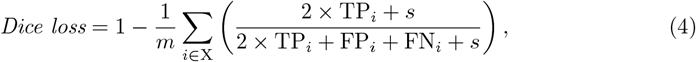

where *s* is a smoothing parameter and *m* is the number of samples in the dataset/mini-batch. FP_*i*_, FN_*i*_, TP_*i*_, and TN_*i*_ denote the number of false positives, false negatives, true positives, and true negatives in the learner’s prediction of sample *i* in X, respectively.

The *Binary Cross entropy loss (BCE)* is given by:

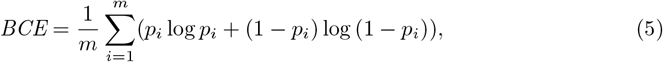

where *m* is the number of samples.

The final *DiceBCE* loss is given as the sum of Eq (4) and Eq (5):

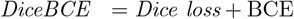

The target learner approximates the optimal posterior distribution of its parameters through variational inference by maximizing the log evidence lower bound. The learner,denoted by ℳ, estimates the output *y* for a given input *x*. The initial parameters (*w*_*µ*_, *w*_*ρ*_) represent the prior Bayesian distribution over the true function ℳ that generated the data. The likelihood of the prediction is given by *p*(*y*|ℳ, *x*). The parameters of ℳ are estimated as *p*(ℳ|*X, Y*), where X and Y are sets of inputs and outputs, respectively. Hence, the likelihood of the data is given by,

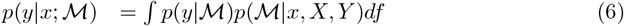

Eq (6) is intractable as an infinite number of parameters can be used to model ℳ. Hence, the parameters of ℳ are approximated by introducing a parameter *w*, which simplifies the problem of finding the best possible parameters.

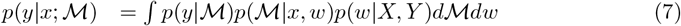

Here, the true distribution *p*(*w* | *X, Y*) remains intractable, and is replaced with a variational distribution *q*(*w*), estimated using variational inference [48,49]. This distribution should be close to the posterior weight distribution of the true function ℳ, given by *p’(w)*. Kullback-Leibler (KL) Divergence [20,30,50,51] measures the distance between two distributions, which estimates the loss between the current and target distributions.

Prior approaches have suggested finding a variational approximation to the Bayesian posterior distribution on the weights [52]. Variational learning finds the parameters *θ* of a distribution on the weights *q(w* |*θ)* that minimizes the KL Divergence of the true Bayesian posterior on the weights.

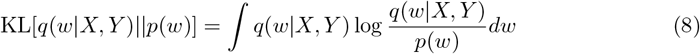

The proposed method does this by sampling weights from the estimated distribution *q*(*w* | *X, Y*) rather than the true posterior *p(w)*. In this approach, where the model parameters are sampled from Gaussian distributions, the KL Divergence can be computed as follows:

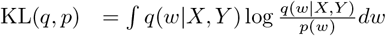

In the case of normal distribution, we can bind the range to (−∞, ∞).

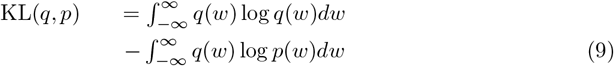

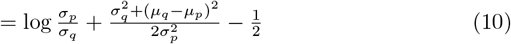

The final loss functions combine the DiceBCE and a scaled KL divergence, resulting in the following:

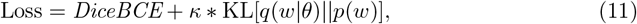

where *κ* is a scaling factor for the KL divergence.

The pipeline we introduce trains the target learner for a given dataset X, Y following Algorithm 1.

#### Algorithm 1

Model Train

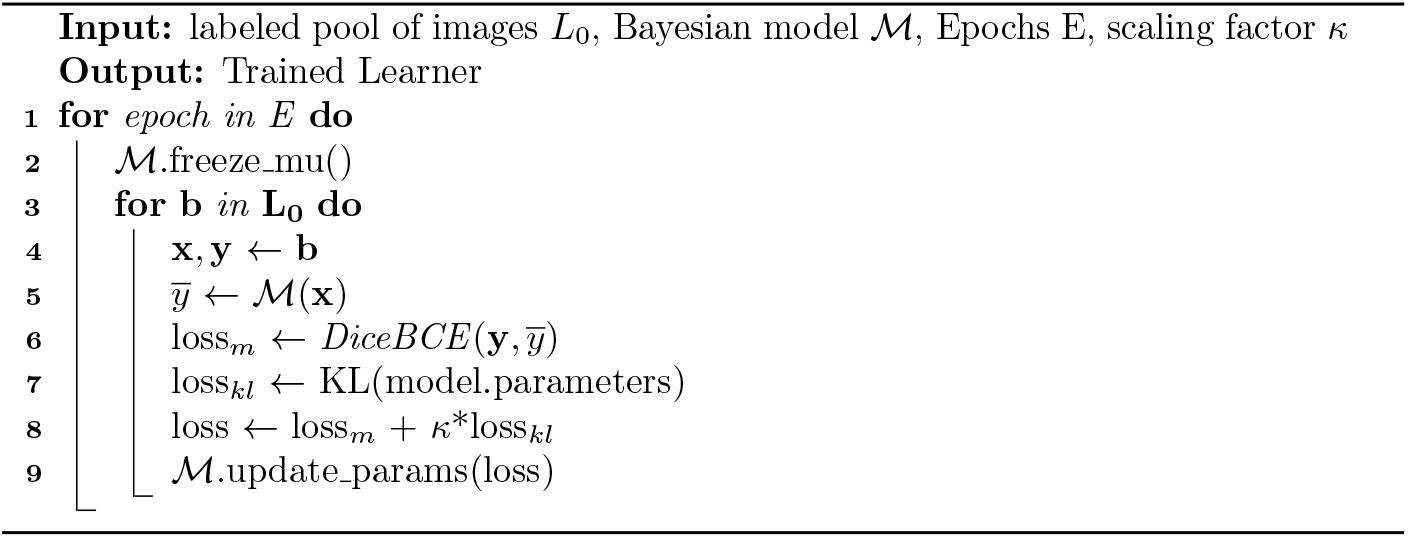

### Estimating Pixelwise Uncertainty

There are two forms of uncertainty: Aleatoric uncertainty - uncertainty due to the inherent ambiguity of the data; and Epistemic uncertainty - uncertainty due to insufficient field knowledge. Aleatoric uncertainty cannot be reduced by adding more data to the training set, as this is a form of uncertainty caused by the inherent nature of the data. In contrast, Epistemic uncertainty can be addressed by including more samples with higher uncertainty. In Bayesian modeling, not only can the uncertainty of a sample be computed, but Aleatoric and Epistemic uncertainty can also be differentiated using the predictive variance of a sample [18,21,22,41].

In classification tasks, finding the total predictive variance to estimate the uncertainty of the sample can be useful. However, current approaches cannot be used directly in image segmentation; localization of uncertainty is not captured in current methods, which is necessary for image segmentation. In most cases, providing an image with high uncertainty to the oracle annotating images is the most useful. As such, sections with high uncertainty are highlighted for manual labeling, hence reducing the overall workload of manual annotation.

The learner’s prediction performance on an unseen sample is computed as the average expectation given the trained learner posterior. Using the equation decomposition in [17], Epistemic uncertainty can be quantified in the image, which is then mapped to its corresponding superpixel. This produces a heatmap of uncertain sections that is then provided to the annotator for labeling. The final labeled image is then added to the labeled pool of images for the next active iteration.

The predictive variance can be decomposed on any new input *x*^*\**^ as:

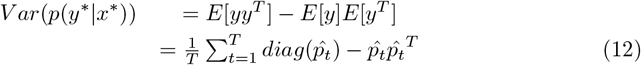

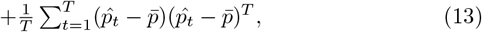

where 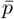 is the average probability of the class, and *T* is the size of the active batch. Eq (12) characterizes the Aleatoric uncertainty and Eq (13) characterizes the Epistemic uncertainty. The method makes use of differentiation between uncertainties caused due to the inherently noisy nature of the data and uncertainty caused due to the learner itself, allowing the learner to accurately find samples in the unlabeled pool of images that carry the most information. The method uses these superpixels to map the computed uncertainties, taking the average divergence of pixel predictions within each superpixel, and in doing so, provides a significant advantage: an objective metric that prioritizes groups of uncertain pixels over single uncertain pixels, maximizing information gain at each active iteration.

The Divergence score for a single superpixel is given by:

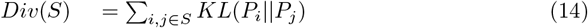

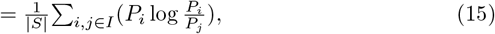

where *S* is a superpixel.

This uncertainty metric uses both forms of uncertainty while evaluating images, fractionally weighting Aleatoric uncertainty to prioritize reducing Epistemic uncertainty. Further, the average superpixel divergence is computed for each unlabeled image while sampling the unlabeled pool of images. When an image is presented to the annotator, the individual superpixels are highlighted based on their respective divergence score, as shown in Fig 1.

**Fig 1.**
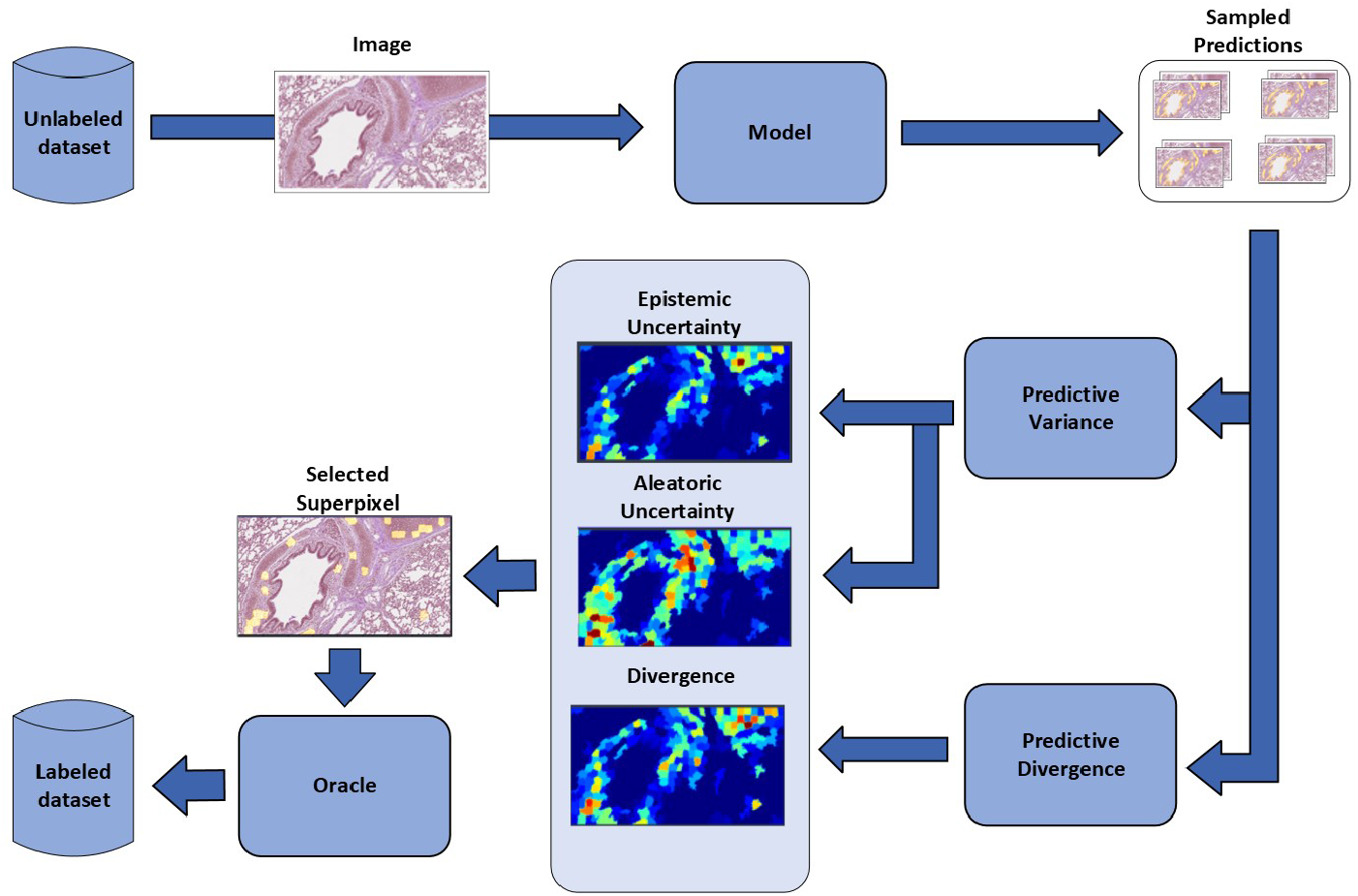
Active Learning Pipeline. Starting from the top left, an image is randomly selected from the pool of unlabeled images. The learner then attempts to segment the image, returning a confidence score on each image pixel. The confidence scores’ Predictive Variance and Predictive Divergence are then computed and used to find the Epistemic uncertainty, Aleatoric uncertainty, and Divergence in prediction. These metrics find the most informative superpixels in the image and provide them to the oracle. The oracle annotates the image and adds the image to the pool of labeled images.

The total uncertainty for an unlabelled sample is computed as an average over *L* independent predictions, as shown in Eq (16)

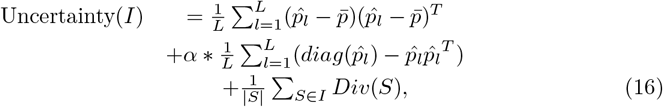

where *α* is a scaling factor for Aleatoric uncertainty, *Div*(*I*) as described in Eq (14), *S* is a superpixel in the input sample, and *I* is an input sample.

### Selecting Regions for Annotation

The regions of uncertainty are often sparsely found throughout a single image. They appear near the boundaries of objects and only in small, dense hotspots. Small sections spread out over a large image make annotating the whole image unnecessarily cumbersome, as relabeling an entire image is slow. Popular approaches tackle this problem by using superpixeling, a clustering heuristic that groups pixels into unique partitions based on local similarity. Using superpixels allows human annotators to minimize labeling costs by annotating the image by uniform regions instead of each individual pixels. We employ SLIC [36] to find the superpixels of an entire image.

The pipeline trains a BNN to perform semantic segmentation on a given corpus following Algorithm 2. Each iteration, defined here as an *active epoch*, begins with the initialization of a BNN, whose starting parameters are fixed for an active training loop and is trained on the most recently updated pool of labeled images. The trained learner then samples the unlabelled pool of images for the most informative samples. The chosen samples are provided to the annotator with clear markings indicating where annotations are required [53], reducing the cost of labeling significantly as annotation is only necessary for highlighted regions.

#### Algorithm 2

SMART

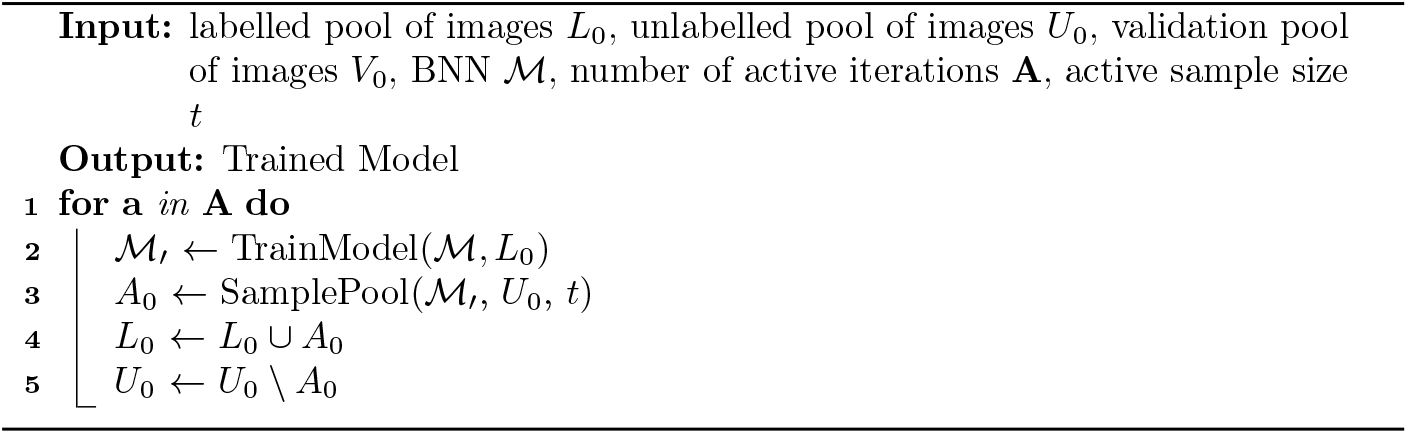

## Results

This section presents the experimental results of the proposed AL paradigm and outlines the testing methodology employed. Section Datasets details the protocols used in the curation of the benchmark datasets and the custom collected pulmonary histology dataset. Subsequently, we describe image augmentation techniques used to introduce more variation on the training data. In Sections Experimental Settings and Comparison Study we describe the computational hardware used to test a model using SmartHisto and SOTA methods it was compared against, and the standardized benchmark datasets used in the comparison, respectively. The source code for the pipeline is available in the project repository at https://github.com/sriram98v/histology_segmentation.

### Experimental Settings

The AL paradigm and other comparative methods were tested under the proposed sampling heuristic with a modified Dice Coefficient loss layer. Images of different sizes were rigidly registered to a reference image size of 256×256. A Bayesian-UNet was trained end-to-end, with cost minimization on 100 epochs performed using an Adam optimizer [54] with an initial learning rate of 0.01. An initial label pool size of five percent was used for the first active epoch, with increments of five percent for each active epoch. The training time for this network was approximately 2 hours on a workstation with NVIDIA GeForce RTX 3090 GPU. The output of the last convolutional layer with sigmoid non-linearity consisted of a probability map for each class. Pixels with computed probabilities of 0.6 or more were considered to belong to the respective output class of the corresponding dataset.

### Datasets

The comparative study uses the publicly available histology dataset GlaS [4] Further, we also include a pulmonary histology dataset used in the ablation study of the proposed approach, as the benchmark datasets are binary segmentation datasets, not multi-class datasets, better reflecting semantic image segmentation in bioinformatics.

#### Glas Dataset

The GlaS dataset included 167 images from 16 H&E stained histological sections of stage T3 or T42 colorectal adenocarcinoma. Each section belonged to a different patient, and sections were processed in the laboratory on different occasions. Thus, the dataset exhibits high inter-subject variability in stain distribution and tissue architecture. The digitization of these histological sections into whole-slide images (WSIs) was accomplished using a Zeiss MIRAX MIDI Slide Scanner with a pixel resolution of 0.465µm.

#### Pulmonary Histopathology Dataset

The Pulmonary Histology dataset included 235 training and 129 testing tissue images, all of resolution 512×512 pixels containing manually annotated structures and cell types using LabelMe [55]. Histologically normal lung tissue originated from clinically healthy, negative control pigs housed at the National Animal Disease Center (NADC) following animal use protocols approved by the NADC. Each image in this dataset was extracted using Aperio ImageScope from one of four individual WSI (scanned at 400x) scanned at a ×40 equivalent resolution on a Leica Aperio AT2. Four individual WSI were considered sufficient due to the low variability observed across histologically normal lung tissue. The labeled data included respiratory epithelium, alveolar septa, cartilage, and smooth muscle. The images were collected from different experiments with and without image color management within Aperio ImageScope, which introduced sources of variation due to the differences in the staining practices and image scanners across slides. Representative 512×512 sub-images from nuclei dense areas were extracted from WSIs to reduce the computational burden of processing multiple WSIs. The dataset includes one crop per WSI and animal to ensure diversity. Both epithelial and stromal nuclei were manually annotated in the 1000×1000 sub-images using Aperio ImageScope. In order to use the WSIs in the proposed pipeline with limited memory, ten sub-images, and their corresponding annotations were randomly sampled from each of the images at a resolution of 250×250 pixels, resulting in 233 training images and 100 testing images.

#### Image Augmentation

Each image was subject to a series of independent random transforms. All samples from the training dataset underwent a set of transforms randomly applied to the image with a probability of 0.5. Variable-degree flips, Gaussian noise, and Gaussian blur were employed to augment the images, applied in series to each image before training. Traditionally, color channels shifts and swaps are also employed in image augmentation. However, we do not employ any form color alterations in image augmentation as such alterations to input images may make the impact of histopathology tissue staining, hindering the performance of the final model.

### Comparison Study

The learner performance on each of the previously listed datasets are presented along with the performance of other benchmark models. The masks predicted by our learner on each of the datasets is displayed in their corresponding subsections. Our method was compared to some benchmark methods such as **ENT, MAR** and **RAND**, and SOTA methods such as **QBC** [37] **and DEAL** [38]. The performance of each of the sampling methods was evaluated as the average performance over 10 full runs of the active sampling method over a full dataset, where for each run, the starting parameters of the learner were drawn from independent and identical distributions.

#### Performance on GlaS Dataset

The comparative results of our method against benchmark methods using the GlaS dataset is plotted in Fig 2. Our learner reaches a better performance than the benchmarks. The learner trained using our sampling approach outperforms the benchmark methods starting from the fifth active epoch, achieving an mIoU of *≈* 0.75 at the end of training. A key behavior to be noted here is that in the Bayesian settings, the learners consistently identify the parts of the image that are always found to be a part of the gland (the outer ring) with both high accuracy and low uncertainty. The inner segment of each gland is highly variable and is detected as such by the learner as an uncertain area of the image. This uncertainty is highlighted in the uncertainty map produced by the learner. Such identification of the variable and consistent patterns in the input images is a common trend seen in all of the following experiments.

**Fig 2.**
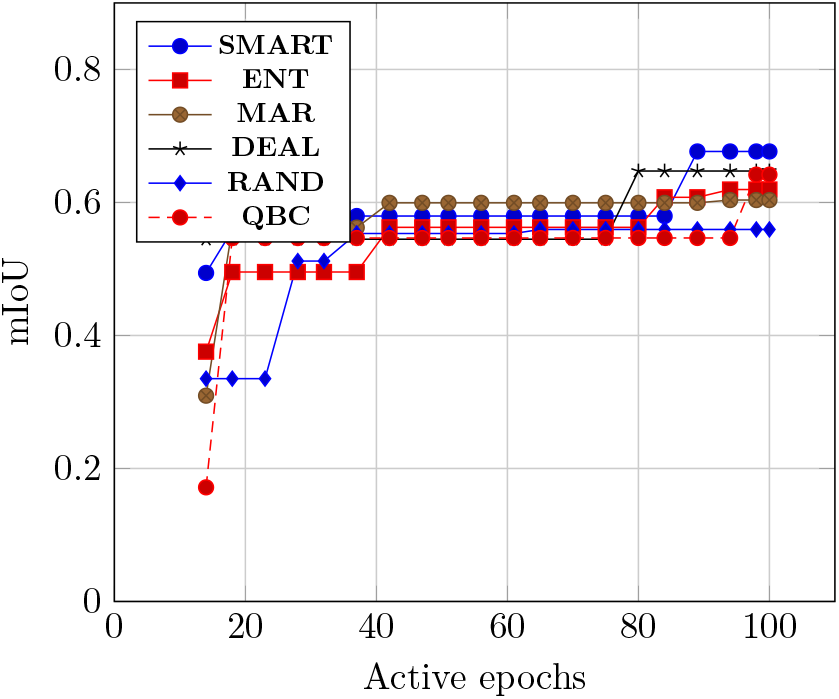
Model Performance after training on each of the sampling methods on the GlaS dataset. Performance of trained using each of the benchmark methods. Our proposed method outperforms the state-of-the-art methods in each consecutive active iteration.

#### Performance on the Pulmonary Histology Dataset

The comparative results of our method against the previously listed benchmark methods is plotted in Fig 3. Our method reaches a better performance than the other benchmarks. The predictions by each of the learners trained with each of the sampling techniques is displayed in Fig 4. The predictions for each target class by each of the methods is listed with their corresponding class name. Fig 5 shows the uncertainty maps computed per label class. Note that there is a high concentration of uncertainty on the border of each class map rather than the center. This is likely due to the variability in annotation accuracy from a human annotator.

**Fig 3.**
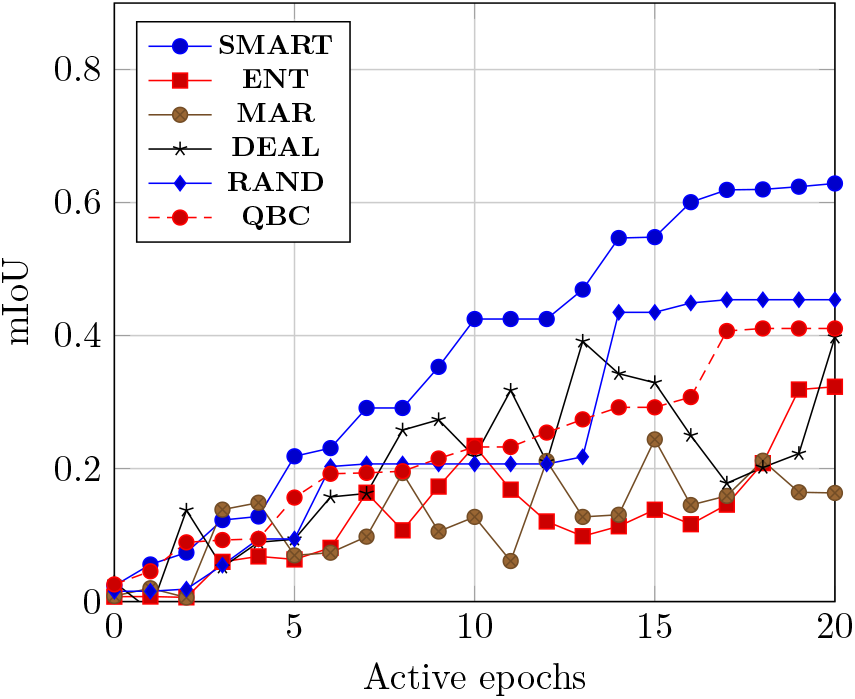
Model Performance after training on each of the sampling methods on the Pulmonary Histology dataset. Performance of trained using each of the benchmark methods. Our proposed method outperforms the state-of-the-art methods in each consecutive active iteration.

**Fig 4.**
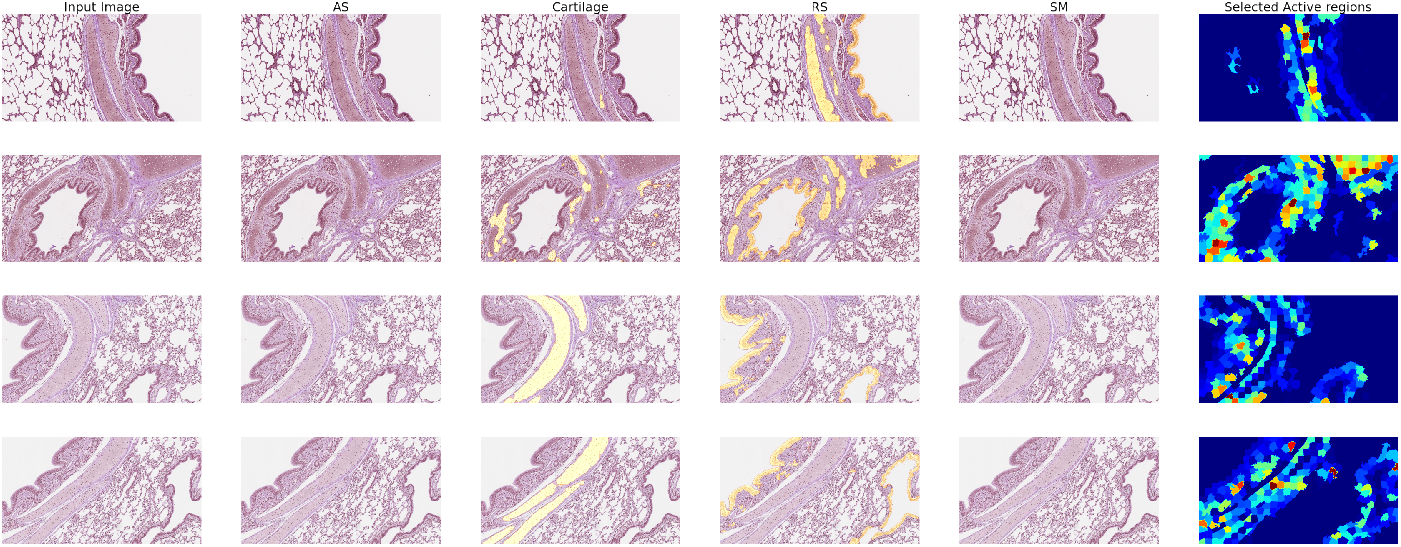
Model Performance and Predictions after training on each of the sampling methods on the Pulmonary Histology dataset. (A) Performance of trained using each of the benchmark methods. Note that our method outperforms the standard benchmark and the state-of-the-art methods methods in each consecutive active iteration. (B) Starting from the leftmost column, each column represents the input image, Ground Truth (GT) binary mask, predicted mask by each of sampling methods as indicated by the column title. Each row indicates the class label of the predicted mask.

**Fig 5.**
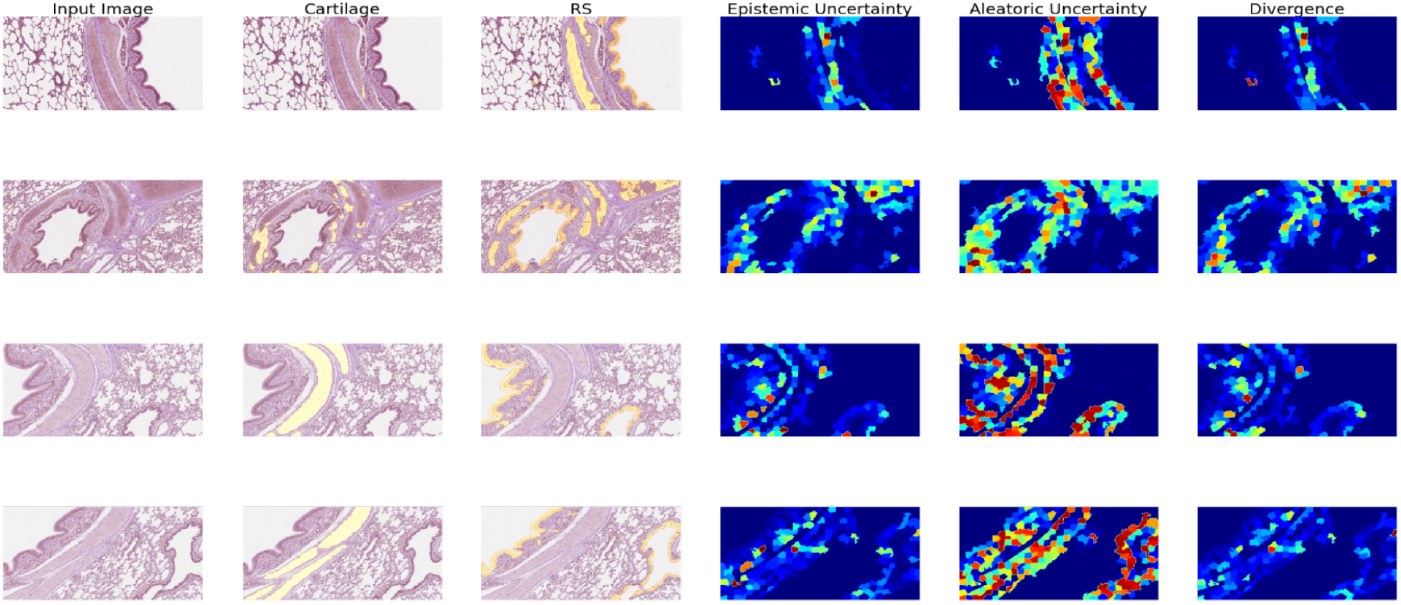
Uncertainty maps computed by the learner trained with the proposed sampling technique on the Pulmonary Histology dataset. The uncertainty maps for predicted class labels as computed by the SmartHisto approach. Note that the borders of each of the classes have a higher uncertainty as they are more variable in each of the input training images.

### Ablation Study

In this section, the results of an ablation study on the parameters of our sampling method using the pulmonary histology dataset described previously in Section Pulmonary Histopathology Dataset are provided with a comparison against the baseline sampling methods **RAND, ENT**, and margin sampling (**MAR**).

#### Effect of KL weight

In order to show the effect of KL Divergence on the overall performance of the learner training within each active epoch, the learner performance was evaluated with a range of values for *κ* in Eq (11) in the loss function. A fixed training set and a distinct validation set are used to evaluate the learner performance and compare the loss of the learner during each training epoch along with the learner accuracy on the validation set. As shown in S1 Fig (a), the learner performances on four different values of *κ ∈ {* 0, 1*}* is plotted at each active epoch, with *κ* = 1 providing the best overall performance.

#### Effect of aleatoric uncertainty weight

In order to evaluate the contribution of aleatoric uncertainty to the active learning sampling scheme, the performance of the learner was evaluated with a range of different values for *α* in Eq (16) of the loss function. Full active training loops were used for each of 4 different values of *α ∈ {* 0, 1*}* without including the divergence term described in Eq (14). The learner performance was evaluated at each active epoch and plotted for each value of *α*. S1 Fig (b) shows that *α* = 1 displays the best overall learner performance.

#### Effect of Divergence

To evaluate the contribution of divergence to the active learning sampling scheme, learner performance with and without the divergence term in Eq (16) were compared while setting *α* = 0. The learner was run once on each setting for a full active learning cycle and was evaluated at the end of each active learning epoch. S1 Fig (c) shows the best learner performance with all the sampling terms.

## Discussion

This paper introduces a novel Active Learning paradigm that uses a Bayesian approach to reduce annotation costs in image segmentation. The approach employs a deep Bayesian active learning method for semantic segmentation,leveraging a Bayesian Neural Network to identify areas of high uncertainty in images. By using an uncertainty metric to segment images, we were able to distinquish when poor classifications were due to deficient model knowledge rather than the inherently noisy nature of the data.

We also developed an acquisition heuristic that reduces labeling effort for large distributed datasets. This approach has significant utility in histopathology, a field challenged by the high cost of accurately annotating training datasets. The method is robust to overfitting and enables faster model convergence through intelligent sample selection. Additionally, SmartHisto is sufficiently flexible to be reapplied to any neural network architecture for image segmentation, substantially reducing the number of samples that require accurate annotations.

We demonstrate consistent performance improvements across all datasets for semantic segmentation. More generally, this framework can be applied to multiple computer vision tasks, including instance segmentation, object localization, and pose estimation.

## Acknowledgments

We sincerely thank Dr. Li for their valuable comments and insightful discussions, which greatly improved this work. This work was supported in part by the USDA-ARS (ARS project numbers 5030-32000-231-000D and 5030-32000-231-095-S); the NIAID, NIH, and HHS (Contract No. 75N93021C00015); and the SCINet project of the USDA-ARS (ARS project number 0500-00093-001-00-D). The funders had no role in study design, data collection and interpretation, or the decision to submit the work for publication. Mention of trade names or commercial products in this article is solely for the purpose of providing specific information and does not imply recommendation or endorsement by the USDA. USDA is an equal opportunity provider and employer.

## Supporting information

**S1 Fig. Ablation study of the proposed method using the Pulmonary Histology dataset**. (a) Ablation study of our loss function. A fixed training set and a distinct validation set were used to evaluate learner performance and compare the loss of the learner during each training epoch along with learner accuracy on the validation set. As shown in the plots, the tested and compared learner performances used four different values of *κ* ∈ {0, 1} which was plotted at each active epoch, with *κ* = 1 providing the best overall performance. (b) Ablation study of our uncertainty metric: the learner’s performance was evaluated with different values for *α* in Eq (16) of the loss function. Full active training loops were used for each of 4 different values of *α ∈ {*0, 1*}* without including the divergence term described in Eq (14). The model performance was evaluated at each active epoch and plotted for each value of *α*. The plot shows that *α* = 1 provides the best performance of the model. (c) Ablation study of divergence: model performance with and without the divergence term in Eq (16) are compared while setting *α* = 0. The model was run once on each setting for a full active learning cycle and was evaluated at the end of each active learning epoch. The plot shows the best overall model performance with all the sampling terms.

